# Excitatory and Inhibitory Effects of Saphenous Nerve Stimulation on Two Different Bladder Conditons: Underactivity and Overactivity

**DOI:** 10.1101/2024.09.21.614283

**Authors:** Chen Li, Zhonghan Zhou, Limin Liao, Xing Li

## Abstract

**Background:** This study aimed at exploring the effects of saphenous nerve stimulation (SNS) on treating both overactive bladder (OAB) and underactive bladder (UAB).

**Method:** In 6 α-chloralose anesthetized cats, bipolar nerve cuff electrodes were implanted on the saphenous nerve and pudendal nerve. UAB was induced by pudendal nerve stimulation (PNS) at 5Hz, 2 threshold (T) and OAB was induced by infusion of 0.25% acetic acid (AA). Multiple cystometrograms (CMGs) were performed to investigate the effects of SNS on pathological bladder at 1Hz and 20Hz, respectively.

**Results:** Application of PNS (5Hz, 2T) induced UAB by significantly increasing the bladder capacity (BC) to 156.3%±9.8% of control level, while combination of PNS and SNS (1Hz, 2T) applied during CMGs normalized the bladder underactivity by significantly reducing the BC to 93.6%±9.5% (P =0.026). Moreover, the BC was reduced to 64.1%±5.4% of control after infusion of AA, and SNS at 20Hz, 6T significantly increased the BC back to 93.4%±6.3% (P =0.005). No post-stimulation effect of SNS was detected at both 1Hz and 20Hz. However, there were no significant changes of contraction amplitude and duration during stimulation.

**Conclusion:** In this study, we confirmed the frequency-dependence of SNS in regulating pathological bladder in cats. It provided experimental evidence for treating both OAB and UAB using SNS in clinic.

## 1 Introduction

Electrical neuromodulation has been proved to be effective in treating lower urinary tract dysfunction (LUTD). The United States Food and Drug Administration (FDA) has approved two approaches: sacral neuromodulation (SNM) and tibial nerve stimulation (TNS)[1]. Plenty of studies have confirmed the benefits of SNM for treating overactive bladder (OAB) and non-obstructive urinary retention (NOUR)[2]. It’s especially effective for women with Folwer’s syndrome, which is one of the characteristics of underactive bladder (UAB)[3]. TNS is a third line therapy for refractory OAB patients who have experienced previous treatment failure with anticholinergic medicines[4]. Moreover, there are accumulating evidence that TNS also has a potential effect for UAB in both experimental and clinical studies[5,6]. Because peripheral nerve stimulation is less invasive and inexpensive, increasing attention has been focused on the application of TNS in clinic.

The TNS is delivered by a percutaneous needle electrode or a surface electrode which is placed 5 cm above the medial malleolus, with the frequency of 20Hz and voltage of equal to or just below the motor threshold (T). However, the voltage required for inhibition of bladder function is 3-6T in anesthetized rats[7,8] and 2-4T in anesthetized cats[9]. This significant disparity between experimental and clinical studies draws great interests. Cadaver studies revealed that the posterior branch of the saphenous nerve (SAFN) is located approximately where the electrode of TNS is placed[10,11]. A finite element analysis of human lower leg revealed that the SAFN is coactivated when TNS is applied at T[12,13]. These results indicated that SAFN plays a vital role in TNS. Further animal experiment confirmed this hypothesis[14,15]. In our previous study, we found that saphenous nerve stimulation (SNS) increased the bladder capacity (BC) at 20Hz, and combination of TNS and SNS can significantly reduce the voltage required for bladder inhibition[16].

Similar to TNS, SNS also exhibited its function in treating UAB. A recent study revealed that SNS at 1Hz can restore the BC and contraction amplitude (AMP) to normal level in an UAB model[17]. Our previous study demonstrated the frequency dependence of SNS in normal bladder condition[16]. When the bladder was infused by normal saline (NS) which activated the non-nociceptive afferent fibers, the bladder function can be inhibited by SNS at 20Hz and activated at 1Hz. However, under pathological conditions of OAB and UAB, whether SNS can exhibits its therapeutic function needs to be validated furtherly. Therefore, we conducted this study to explore the function of SAFN in pathological models, and aimed to provided experimental evidence for treating both OAB and UAB using SNS in clinic.

## 2 METHODS

### 2.1 Surgical Procedures

A total of six adult cats (male, 6-12 months old, weighting 1.74-3.25kg) were involved in this study. The National Institutes of Health guide for the care and use of Laboratory animals was followed, and the experiment was approved the Animal Care and Use Committee at the Capital Medical University (AEEI-2024-213). The surgical procedures were elaborated in the previous studies (Fig. 1)[16,17]. Briefly, anesthes The National Institutes of Health guide for the care and use of Laboratory animals was followed, and the experiment was approved by the Animal Care and Use Committee at the Capital Medical University. ia was induced by isoflurane (2-5% in oxygen) during surgery and maintained by α-chloralose (initial 65mg/kg and supplemented as needed, through left cephalic vein) during data acquisition. The blood oxygen and heart rate were monitored during the experiment. A middle abdominal incision was made, and a double lumen catheter was inserted into bladder through a small cut on the proximal urethra. One lumen was connected to a pump for bladder infusion, the other was connected to a bladder transducer (MP150; BIOPAC System, Inc., Camino Goleta, CA, USA) for pressure measurement. The ureters were isolated, tied, and cut for external drainage. The left SAFN was exposed via a skin incision on the medial thigh slightly above the knee joint. The right pudendal nerve was isolated through an incision in the region of the sciatic notch. Two custom-fabricated bipolar nerve cuff electrodes were implanted on the SAFN and pudendal nerve, respectively. The stimulation was released through an external stimulus generator (Master-8; AMPI, Jerusalem, Israel). After the surgery, the incisions were closed with sutures.

**Fig. 1.**
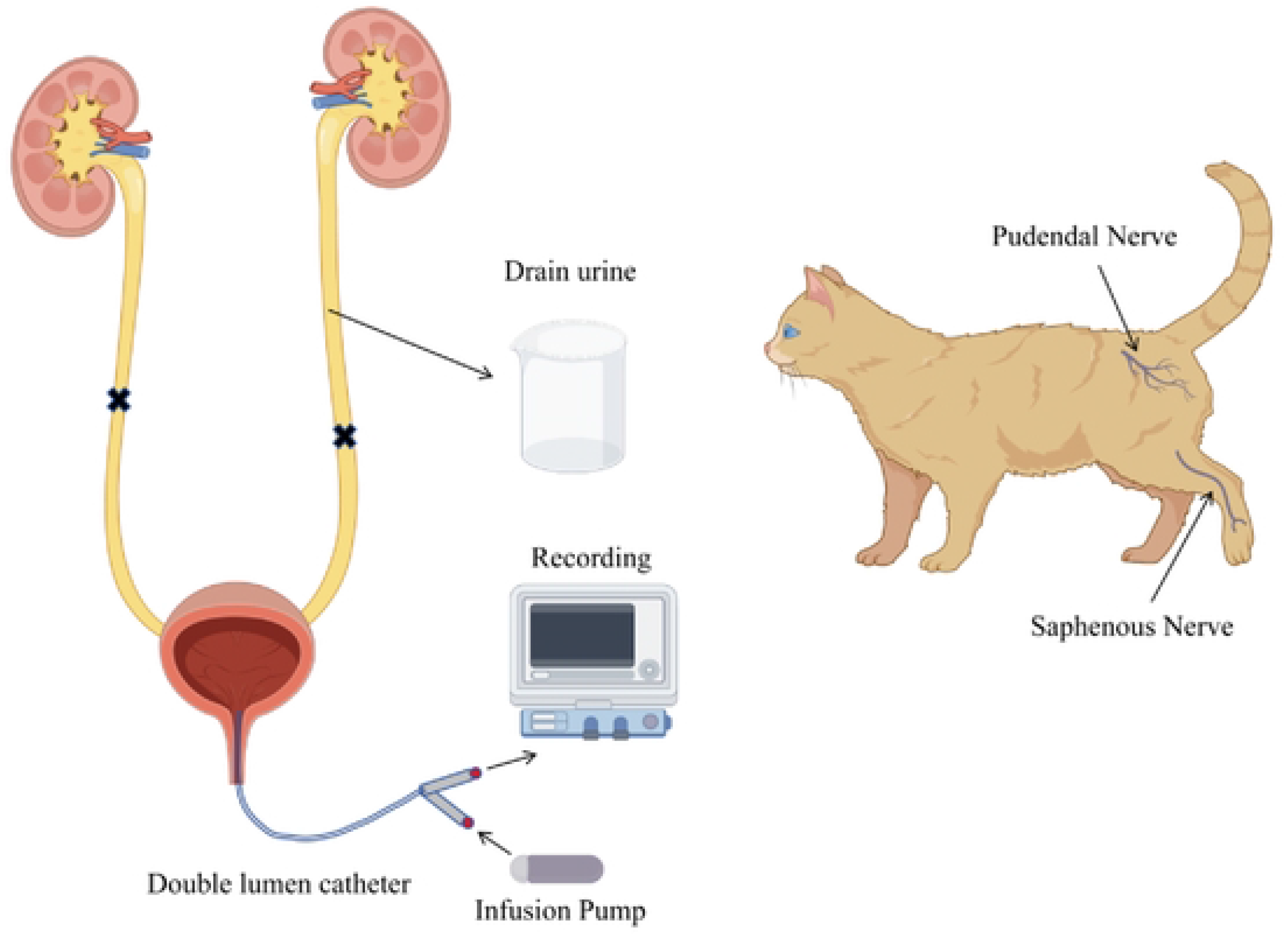
Schematic diagram of the experiment

### 2.2 Stimulation Protocol

Multiple cystometrograms (CMGs) were performed by infusing NS at the rate of 1-2ml/min approximately 30 minutes after the surgery. The BC were defined as the volume threshold for inducing the first micturition reflex contraction (AMP >30cmH2O, contract duration [DUR] >20s). Once the BC was stable, another two or three CMGs were recorded to obtain the baseline of the BC. Afterwards, uniphasic rectangular pulses (0.2ms width) were applied to the nerves. T was defined as the minimal voltage for inducing muscle twitches of the posterior thigh, hip or toe for SAFN, and the anal twitch for pudendal nerve. The whole experiment was divided into two parts. The first section was to investigate the excitatory function of SANF in an UAB model, which was induced by pudendal nerve stimulation (PNS) to mimicking the pathology of Fowler’s syndrome as previously described[18]. 6 CMGs were performed: (1) control CMG without stimulation; (2) CMG with SNS (1Hz, 2T); (3) CMG with PNS (5Hz, 2T), to induce an UAB model; (4) CMG with combination of SNS (1Hz, 2T) and PNS (5Hz, 2T); (5) CMG with PNS (5Hz, 2T), to detect the occurrence of post-stimulation effect; (6) control CMG without stimulation again. The second section was to investigate the inhibitory function of SANF in an OAB model, which is induced by irritation of 0.25% acetic acid (AA). 4 CMGs were performed: (1) control CMG with NS infusion; (2) CMG with AA infusion, to induce an OAB model; (3) CMG with SNS (20Hz, 6T); (4) CMG with AA infusion again, to detect the occurrence of post-stimulation effect. The bladder was evacuated and allowed to rest for 5 mins after each CMG.

### 2.3 Statistical Analysis

Statistical analysis was performed using SPSS version 19.0 software (IBM Corporation, Armonk, NY, USA) and R version 3.5.2 (The R Foundation, Vienna, Austria). Parameters of CMGs, including the BC, AMP and DUA, were measured and normalized to the measurement of the first control result. The data from different animals were presented as mean±standard error. Significant difference (P <0.05) was determined by repeat-measures one-way ANOVA followed by Bonferroni post hoc test.

## 3 RESULTS

### 3.1 The excitatory function of SNS (1Hz, 2T) in the UAB model

SNS at 1Hz, 2T significantly decreased the BC to 64.7%±5.2% of the NS control level (P =0.016). Afterward, application of PNS at 5Hz, 2T increased the BC to 156.3%±9.8% (P =0.033), producing an UAB model characterized by large BC. Then, the combination of SNS and PNS applied during CMG normalized the bladder underactivity by significantly reducing the BC to 93.6%±9.5% of control (P =0.026). The BC returned to the pre-stimulation level, indicating no post-stimulation effect was detected. However, there were no significant changes of the AMP and DUR during SNS or PNS (Figs. 2, 3).

**Fig. 2.**
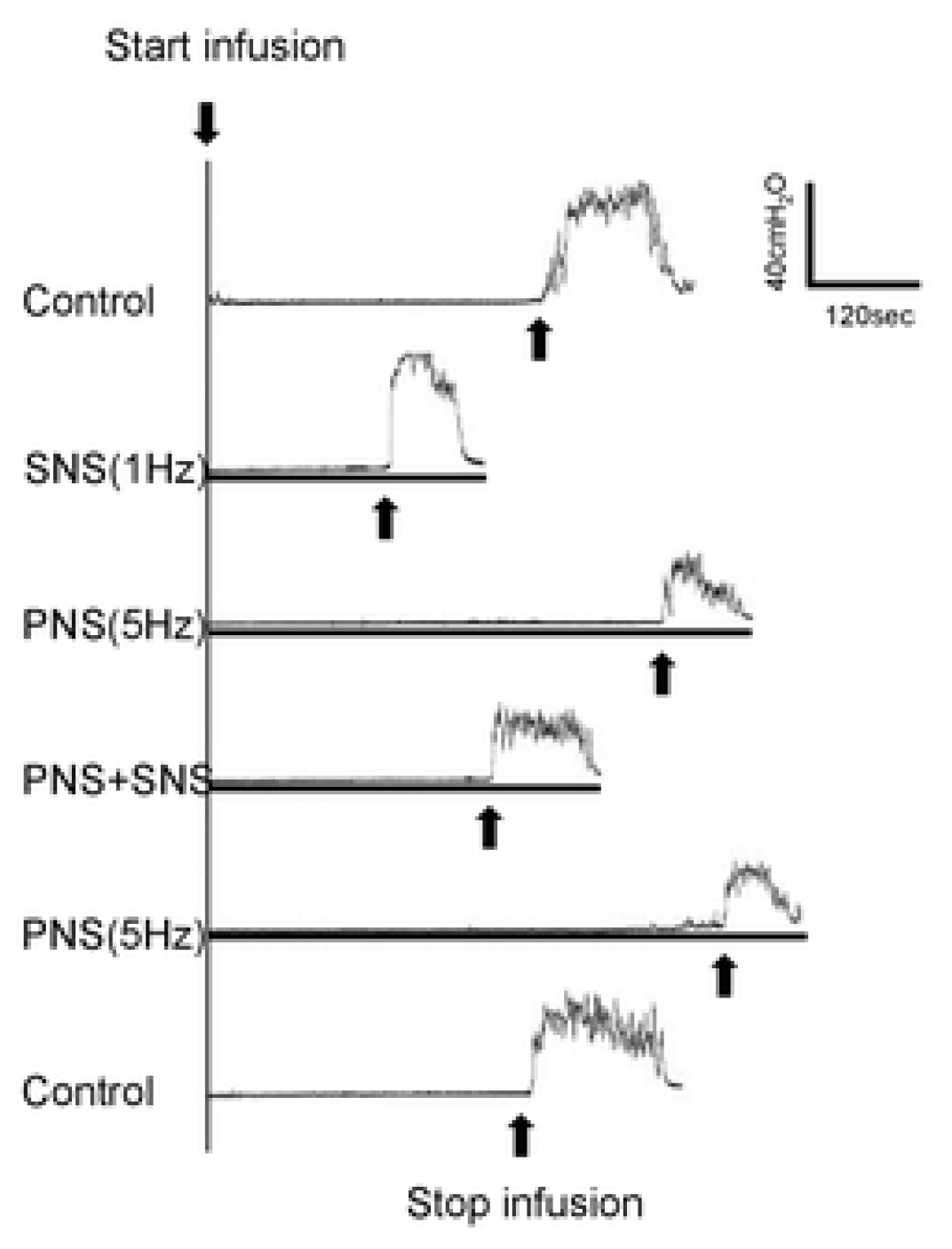
Saphenous nerve stimulation (SNS) at I Hz, 2T applied during a cystometrogram (CMG) normalized the bladder underactivity by significantly reducing the bladder capacity. The black bar under bladder pressure trace represented the duration of stimulation. SNS: IHz, 0.2ms, 2T=0.3V; PNS: 5Hz, 0.2ms, 2T=l.2V; Infusion rate: 1ml/min.

**Fig. 3.**
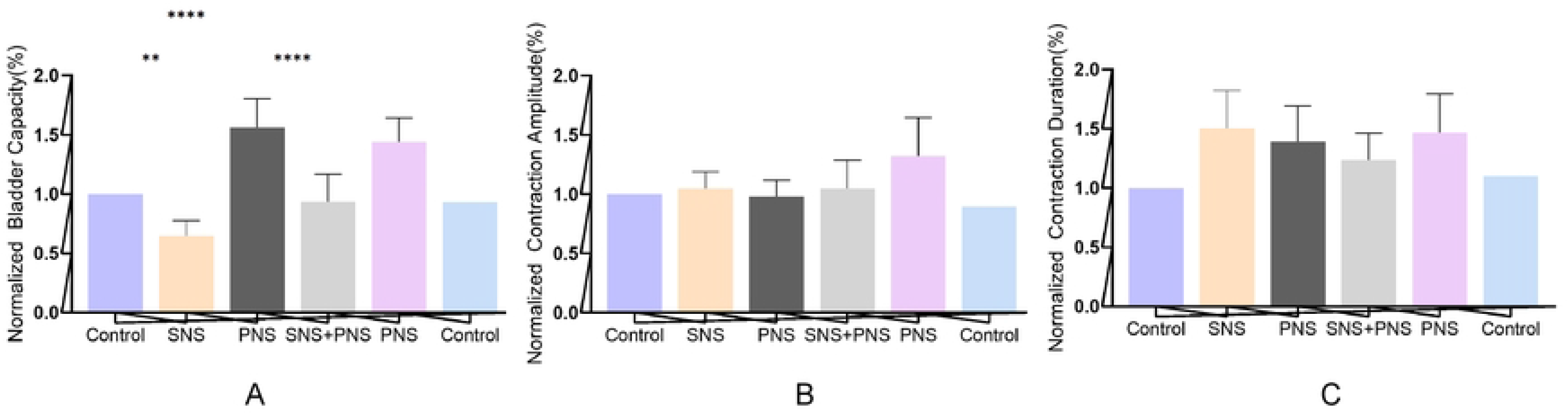
Saphenous nerve stimulation (SNS) at I Hz, 2T on the bladder capacity (A), contraction amplitude (B), and contrac1ion dura1ion (C). SNS: IHz., 0.2ms, 2T=0.3-6.0V; TNS: 5Hz, 0.2ms, 2T=l.2-3.2V; Infusion ra1e: 1-21111/min. *Significantly ditforent (P <0.05, repeat-measures one-way ANOVA followed by Bonferroni post hoc test). N=6 cats.

### 3.1 The inhibitory function of SNS (20Hz, 6T) in the OAB model

The BC was reduced to 64.1%±5.4% of NS control after infusion of AA (P =0.042). Then SNS at 20Hz, 6T significantly increased the BC to 93.4%±6.3% (P =0.005). Similarly, there was no post-stimulation effect on the BC. In addition, the AMP and DUR was not significantly changed by either AA irritation or SNS at 20Hz, 6T (Figs. 4, 5).

**Fig. 4.**
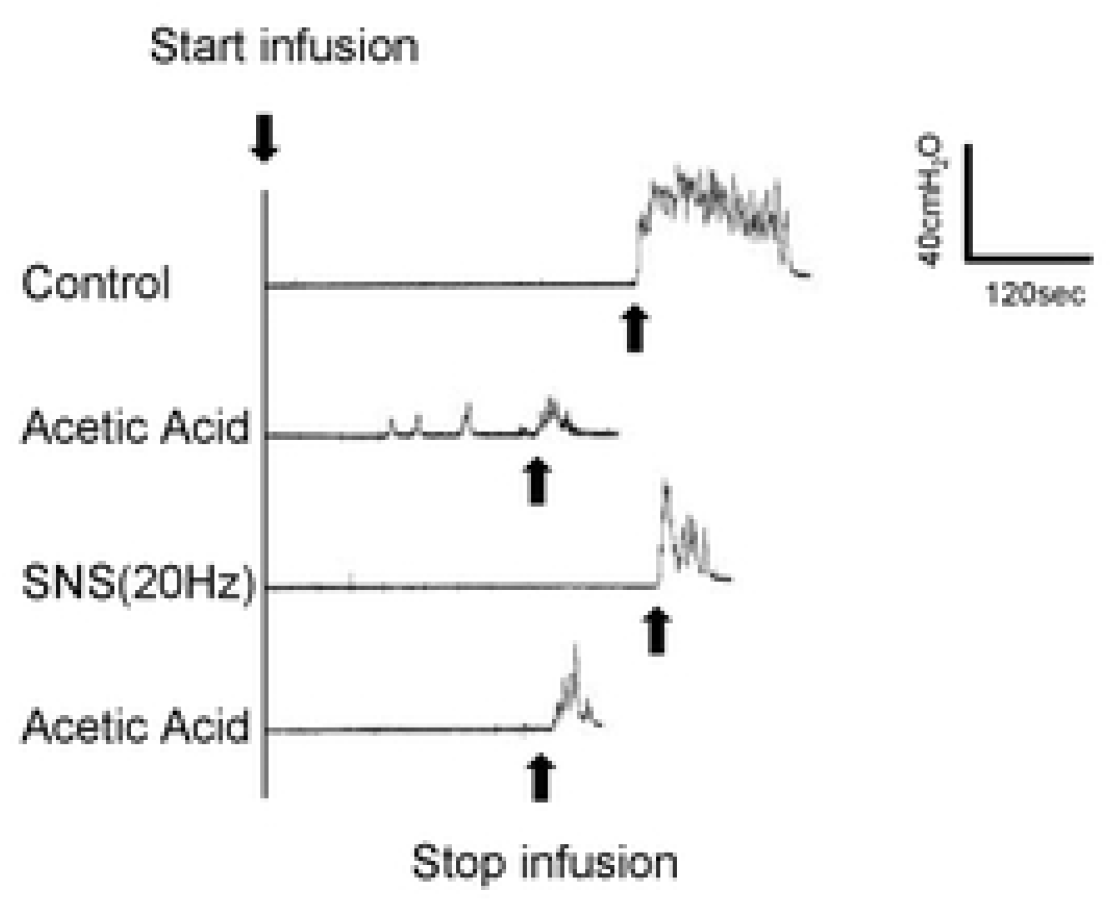
Saphenous nerve stimulation (SNS) at 20Hz, 6T applied during a cystometrogram (CMG) significantly inhibited the irritated bladder. The black bar under bladder pressure trace represented the duration of stimulation. SNS: 20Hz, 0.2ms, 6T=l.2V: Infusion rate: 1ml/min.

**Fig. 5.**
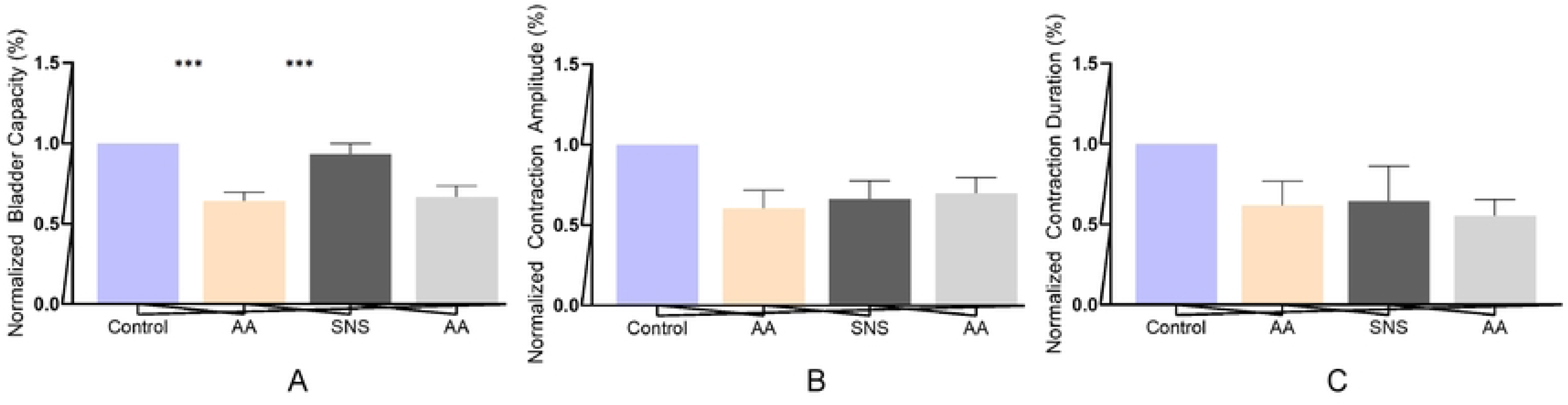
Saphenous nerve stimulation (SNS) at 20Hz, 6T on the bladder capacity (A), contraction amplitude (B), and contraction duration (C). SNS: 20Hz, 0.2ms, 6T=l.2-7.8Y; Infusion rate: 1-2ml/min. *Significantly different (P <0.05, repeat-measures one-way ANOYAfollowed by Bonferroni post hoc test). N=4 cats.

## 4 DISCUSSION

There were several previous studies investigating the role of SAFN in bladder control. Moazzam et al. reported that SNS at 20Hz can increase the BC with NS infusion in anesthetized rats, however, SNS at 2Hz did not have a significant effect on bladder[14,15]. Li et al. reported that when bladder was infused with NS in cats, SNS at 1Hz didn’t change the BC and AMP, while the DUR was significantly increased compared to control[17]. However, SNS at 1Hz normalized bladder underactivity induced by repeated TNS, showing significant difference in the BC, AMP and DUR. But in this study, changes in bladder activity were not detected when SNS was applied at 20Hz. In our previous study, we demonstrated that SNS at 1Hz, 1T or 2T can reduce the BC, while SNS at 20Hz, 6T can significantly increase the BC[16]. This inconsistence may be caused by distinct bladder conditions (physiological or pathological), the different neural circuits of the two species (rodents or cats), various anesthesia (urethane or chloralose) and diverse voltage used in different studies. In this study, we confirmed the frequency-dependence of SAFN in pathological bladder. SNS at 1Hz, 2T can normalized the bladder underactivity, while SNS at 20Hz, 6T can inhibit irritated bladder. The frequency dependence of SAFN is consistent with other peripheral nerves, including TNS (excitatory: 1-2Hz; inhibitory: 5-20Hz), pudendal nerve (excitatory: 3Hz; inhibitory: 20Hz) and sacral dorsal root ganglion (excitatory: 0.25-1.5Hz and 15-30Hz; inhibitory: 3-7Hz)[5,19-21]. In addition, it’s interesting to find a similar pattern of both excitatory and inhibitory frequency between TNS and SNS.

Considering the superficial location of SAFN, it’s reasonable to conduct percutaneous SNS similar to TNS. In a pilot clinical study, a total of 18 OAB patients received SNS at 20Hz, and 14 (87.5%) achieved positive response. Our study furtherly provided experimental evidence for treating OAB with SNS at 20Hz. Previous studies revealed that coactivation of SNS is a potential therapeutic mechanism of TNS, and a combination of SNS and TNS at 20Hz can enhance the inhibitory effects on bladder activity[16]. Therefore, it’s feasible to investigate the combined SNS and TNS for treating OAB patients in the future.

UAB remains to be a great challenge for clinicians. Currently, there was no medical therapy approved by FDA. In comparation to OAB, there are fewer basic and clinic research focused on mechanisms and treatment of UAB. Recently, a study revealed a potential role of TNS at 1Hz in normalizing the bladder underactivity[5]. Similarly, we showed the therapeutic effect of SNS at 1Hz. Thus, it’s possible to explore the application of TNS, SNS, or the combination of TNS and SNS in UAB in clinic.

Recently, Franz et al. tried to investigate the relation between SAFN and hypogastric nerve, which can inhibit bladder function by activating bladder neck and urethra, and relaxing detrusor muscles[22]. However, they found that hypogastric nerve did not mediate the inhibitory effect of SNS. It’s hypothesized that SNS can be coactivated when TNS is applied, but the exact underlying mechanism is still confusing. Therefore, the mechanism of SNS and the relation between SNS and TNS must be explored in further studies.

## 5 CONCLUSION

In summary, we confirmed the frequency-dependence of SAFN in regulating pathological bladder in cats. It provided experimental evidence for treating OAB or UAB with SNS at 20Hz or 1Hz, respectively. Additional clinical studies are required to investigate the effectiveness of SNS or the combination of TNS and SNS. The mechanism of SNS still remains to be explored.

## LIST OF ABBREVIATIONS

LUTD: lower urinary tract dysfunction
FDA: Food and Drug Administration
SNM: sacral neuromodulation
TNS: tibial nerve stimulation
OAB: overactive bladder
NOUR: non-obstructive urinary retention
UAB: underactive bladder
T: threshold
SAFN: saphenous nerve
SNS: saphenous nerve stimulation
BC: bladder capacity
AMP: contraction amplitude
NS: normal saline
CMGs: cystometrograms
PNS: pudendal nerve stimulation

## Statement

The experiment was approved by the Animal Care and Use Committee at the Capital Medical University (AEEI-2024-213)

## Funding

This study was funded by the Beijing Natural Science Foundation (No. 7222235) and the Research Projects of China Rehabilitation Research Centre (2021zx-11).

## Acknowledgement

Figure1 support was provided by Figdraw

## Competing interests

The authors declare that they have no competing interests.

## Notes

### Competing Interest Statement

The authors have declared no competing interest.

